# Accounting for tumor heterogeneity using a sample-specific error model improves sensitivity and specificity in mutation calling for sequencing data

**DOI:** 10.1101/055467

**Authors:** Yu Fan, Xi Liu, Hughes Daniel S. T., Jianjua Zhang, Jianhua Zhang, P. Andrew Futreal, David A. Wheeler, Wang Wenyi

## Abstract

Subclonal mutations reveal important features of the genetic architecture of tumors. However, accurate detection of mutations in genetically heterogeneous tumor cell populations using NGS remains challenging. We developed MuSE (http://bioinformatics.mdanderson.org/main/MuSE), mutation calling using a Markov substitution model for evolution, a novel approach modeling the evolution of the allelic composition of the tumor and normal tissue at each reference base. MuSE adopts a sample-specific error model that reflects the underlying tumor heterogeneity to greatly improve overall accuracy. We demonstrate the accuracy of MuSE in calling subclonal mutations in the context of large-scale tumor sequencing projects using whole exome and whole genome sequence.

## Background

The detection of somatic point mutations is a key component of cancer genomic research, which has been rapidly developing since next-generation sequencing (NGS) technology revealed its potential for describing genetic alterations in cancer [1, 2, 3, 4, 5, 6]. As the cost of NGS has decreased, the need to thoroughly interrogate the cancer genome has spurred the migration from using whole exome sequencing (WES) to whole genome sequencing (WGS). A critical challenge accompanying this migration is the rigorous requirement of specificity, considering that a false positive rate (FPR) of even 1 per megabase pair (Mbp) results in 3,000 incorrect variant calls for WGS data. In addition, the sequencing depth decreases from 100 − 200× for WES data to 30 − 60× for WGS data, resulting in a lower signal-to-noise ratio and making accurate mutation calling more difficult.

Another nontrivial difficulty is accounting for the influence of tumor heterogeneity that is commonly observed in mutation calling. The presence of both normal cells and tumor subclones in the sample causes this phenomenon to vary from sample to sample [7, 8]. It is thus important to identify sample-specific cutoffs dynamically and report tier-based variant call sets instead of using a fixed cutoff for all samples which is current common practice. On the other hand, tier-based variant call sets that inherently attach uncertainties will be helpful when evaluating the behavior of low VAF mutations and seeking to understand the effect of tumor heterogeneity.

Here, we present a novel and automatic approach to discovering somatic mutations, <u>Mu</u>tation calling using a Markov <u>S</u>ubstitution model for <u>E</u>volution (MuSE), which models the evolution of the reference allele to the allelic composition of the tumor and normal tissue at each genomic locus. We further adopt a sample-specific error model to identify cutoffs, reflecting the variation in tumor heterogeneity among samples. We demonstrate the reliable performance of MuSE, a good balance of sensitivity and specificity, with various types of data.

## Results and discussion

### MuSE design

MuSE comprises two steps (Fig. 1). The first step,‘MuSE call’ (Fig. 1a,b), takes as input the binary sequence alignment map (BAM) formatted sequence data that require special preparation from the pair of tumor and normal DNA samples. The results of our investigation favored the co-local realignment of tumor and matched-normal BAMs rather than the local realignment of tumor and matched-normal BAMs separately (data not shown). MuSE carries out pre-filtering on every genomic locus, which is a common practice [e.g., see 5] ahead of variant detection in order to accelerate the computing speed and remove potential false positives. Next, MuSE accomplishes variant detection by employing the F81 Markov substitution model [9], which provides the estimates of equilibrium frequencies for all four alleles (*π_A_*, *π_C_*, *π_G_*, *π_T_*), and the evolutionary distance (*ν*). In practice, we report the maximum *a posteriori* (MAP) estimates of ***π*** and *ν* instead of exploring the full posterior distribution.

**Figure 1.**
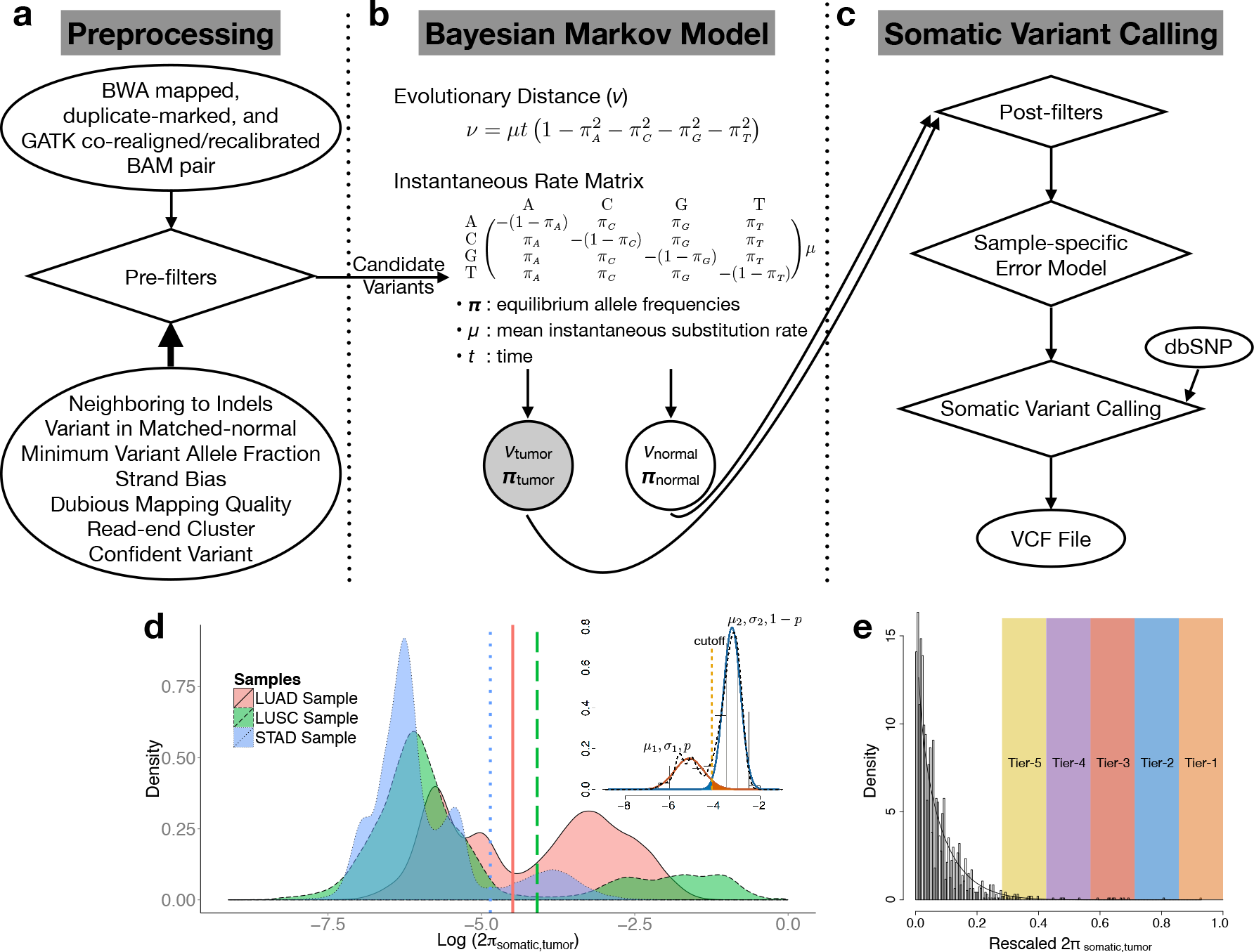
Flowchart of the somatic point mutation caller MuSE. (a) MuSE takes as input the BWA-aligned BAM sequence data from the pair of tumor and normal DNA samples. The BAM sequence data are processed by following the Genome Analysis Toolkit (GATK) Best Practices. Next, at each genomic locus, MuSE applies seven heuristic pre-filters to screen out false positives resulting from correlated sequencing artifacts. (b) MuSE employs the F81 Markov substitution model of DNA sequence evolution to describe the evolution from the reference allele to the tumor and the normal allelic composition. It writes to an output file the MAP estimates of four allele equilibrium frequencies (*π*) and the evolutionary distance (*ν*). (c) MuSE uses the MAP estimates of *π* to compute the tier-based cutoffs by building a sample-specific error model. MuSE deploys two different methods of building the sample-specific error model for the respective WES data and WGS data. Besides using the sample-specific error model, MuSE takes into account the dbSNP information by requiring a more stringent cutoff for a dbSNP position than for a non-dbSNP position. The final output is a VCF file that lists all the identified somatic variants. (d) Illustration of the sample-specific error model for WGS data. Tumor heterogeneity is illustrated using TCGA lung adenocarcinoma (LUAD), lung squamous cell carcinoma (LUSC) and stomach adenocarcinoma (STAD) WGS data. All *π*_somatic_ selected for building the sample-specific error models are used to draw the densities that are on the logarithmic scale. At the top right, we show a two-component Gaussian mixture distribution with means *μ*_1_ and *μ*_2_, standard deviations *σ*_1_ and *σ*_2_, and weights *p* and 1 − *p*, for true negative and true positive, respectively. The expected false positive probability caused by the identified cutoff is the area labelled in red (on the right side of the cutoff), and the false negative probability is the area labelled in blue (on the left side of the cutoff). We first identify a cutoff that minimizes the sum of the two probabilities and add tiered cutoffs that are less stringent than the first one. (e) Illustration of the sample-specific error model for WES data. Selected *π*_somatic_ are rescaled to fit a Beta distribution. Tiers 1 to 5 are labelled for illustration purposes, but not in equal proportion to those in the real data.

The second step, ‘MuSE sump’ (Fig. 1c), takes as input the post-filtered *π*_somatic,tumor_, and computes tier-based cutoffs from a sample-specific error model. As a unique feature of MuSE, the tier-based cutoffs (PASS, Tiers 1 to 5) address the large variations observed in the distributions of *π*_somatic,tumor_ across tumor samples (Fig. 1d). With WGS data, we fit a two-component Gaussian mixture model to log (2*π*_somatic,tumor_) across positions in order to separate true positive mutations from the reference positions, which represent two major modes in the distribution of log (2*π*_somatic,tumor_). However, the degree of difference between the true negative (reference) and true positive (somatic mutation) positions and whether it is detectable depend on the sequencing depth and the VAFs of the mutations. When true mutations are of low VAF, presenting a distribution that largely overlaps with that of the true negative positions, we use a cutoff of 0.005 as our lowest boundary (Tier–5) to control the number of false positives. When the number of true positives is relatively minimal compared to that of true negatives, as in most WES data [mutation rate up to 10/Mbp; 10], we model *π*_somatic,tumor_ as a Beta distribution (Fig. 1e) and call mutations as the extreme and rare events on the right tail of the fitted distribution. We take into account the dbSNP information by requiring a cutoff that is two times more stringent for a dbSNP position than for a non-dbSNP position. The final output of the second step is a variant call format (VCF) file that lists the identified somatic variants.

### Synthetic data

We measured the performance of MuSE using synthetic data and compared sensitivity and specificity of MuSE with that of other state-of-art callers [1, 4, 5, 11]. MuSE is intended to run with little or no human curation. For that reason, all callers were evaluated without human curation to yield a uniform comparison, although in practice, output from mutation callers is often curated. We first made the comparison using the
synthetic data IS1, IS2 and IS3 (9.11 gigabase pair (Gbp)) from the ICGC-TCGA DREAM Mutation Calling challenge [6]. The complexity of the three data sets increased because of elevating mutation rates, declining VAFs and incorporating multiple subclones. This increased data complexity affected the performance of all callers, which was evident in the synchronized decreases in sensitivity Fig. 2a). In all three data sets, MuSE was more sensitive and specific than MuTect, SomaticSniper, Strelka and VarScan2. Moreover, MuSE identified cutoffs varying by the sample (Fig. 2a, bottom right). These cutoffs at the PASS level are located at the top left corners of the receiver operating characteristic (ROC) curves, which suggests an ideal balance between sensitivity and specificity. Since IS1 was the least complex and furthest away from real data, additional tiers were not able to improve sensitivity.

**Figure 2.**
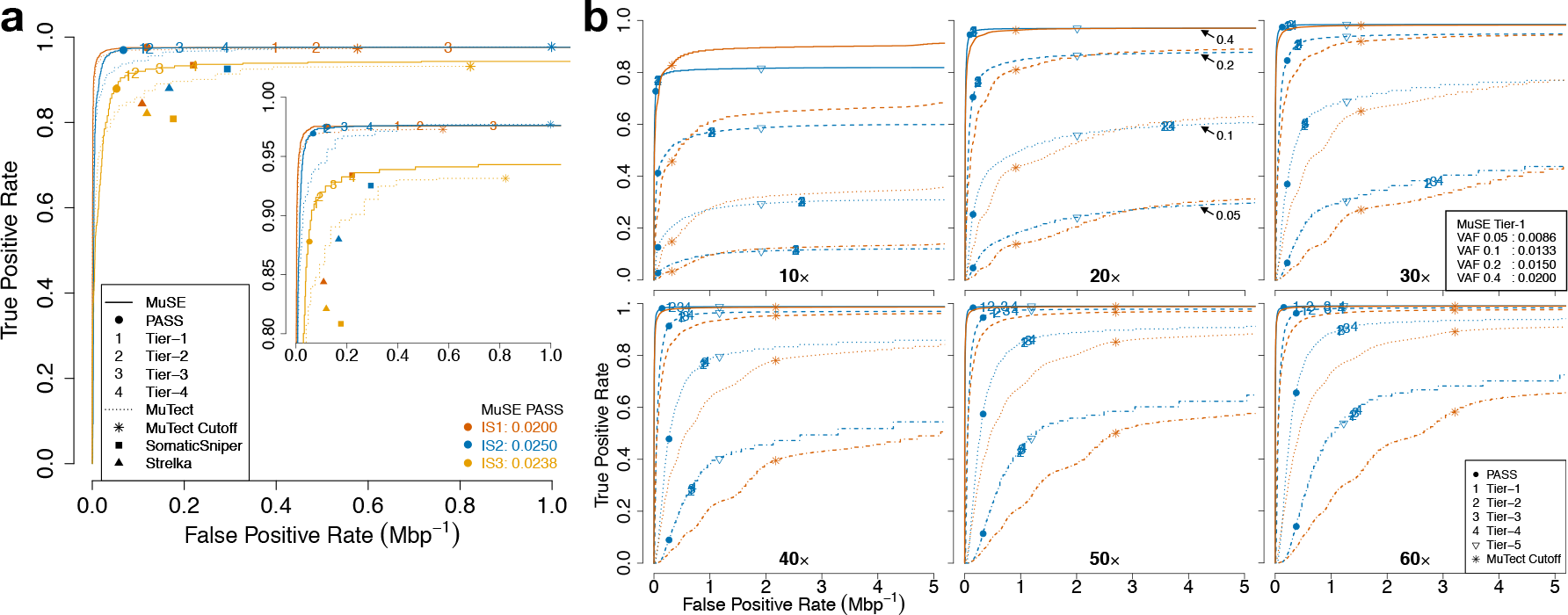
Comparison of sensitivity and specificity of MuSE and MuTect using synthetic data. (a) Comparison of sensitivity and specificity of MuSE (solid line), MuTect (dotted line), SomaticSniper (solid square) and Strelka (solid triangle) using the synthetic data IS1, IS2 and IS3 from the ICGC-TCGA DREAM Mutation Calling challenge. The numbers of positions with positive conditions are 3,535, 4,322 and 7,903, respectively. Both tumor and matched-normal data have ~30× average coverage. The three synthetic data sets are color-coded using red, blue and orange, respectively, and the associated ROC curves, focusing on an FPR between 0 and 1 × 10^−6^, are ordered from top left to bottom right. The tier-based sample-specific cutoffs of MuSE and the MuTect default cutoff are labelled correspondingly. The embedded plot focuses on a narrow range of TPR. The two times when PASS cutoffs were identified are listed at the bottom right corner. Sensitivity and specificity of VarScan2 (not plotted due to out of bounds) were 0.9859 and 8.369 × 10^−6^ (IS1), 0.9704 and 1.294 × 10^−6^ (IS2) and 0.8602 and 1.478 × 10^−6^ (IS3), respectively. (b) Comparison of sensitivity and specificity of MuSE (blue line) and MuTect (red line) using the virtual-tumor benchmarking approach. The ROC curves focus on an FPR between 0 and 5 × 10^−6^. Tumor sample sequencing depth varies from 10× to 60×, and matched-normal sample sequencing depth is fixed at 30×. Four scenarios of spike-in VAF 0.05 (dot-dashed), 0.1 (dotted), 0.2 (dashed) and 0.4 (solid) are plotted for every sequencing depth. The tier-based sample-specific cutoffs of MuSE and the MuTect default cutoff are labelled accordingly. Some MuSE cutoffs are close to each other and overlap on the plot. For 30× coverage, the two times that Tier–1 cutoffs were identified are listed at the bottom right corner of the corresponding subplot.

Furthermore, using the virtual-tumor benchmarking approach [5], we studied the impact of sequencing depth (10× to 60×) and VAFs (0.05, 0.1, 0.2 and 0.4) on MuSE and MuTect in whole genomes (18.2 Gbp; Fig. 2b, Supplementary Table 1). From moderate (30×) to high (60×) coverage, the MuSE curves stayed on top of the MuTect curves. At low (10× and 20×) coverage, the two curves crossed as FPR increased. These two low coverage data sets had low signal-to-noise ratios and were most sensitive to losing true positives from post-filtering. Nevertheless, for the segment of the curve that contained the MuTect default cutoff, the MuSE curve was still on top of its counterpart, except for one scenario, 10× and VAF of 0.4. The incremental changes in calling accuracy from Tier–1 to Tier–4 were more evident in scenarios with high VAFs than in those with low VAFs. Different from the DREAM challenge data, in this data set, Tier–1 cutoffs showed the biggest improvement in sensitivity compared to one level up, PASS, and moved closer to the top left corners of the ROC curves in all simulation scenarios from 30× to 60× coverage, except for 30× and VAF of 0.05. For different VAF spike-in scenarios, again, MuSE identified Tier–1 cutoffs that were distinct from each other (Fig. 2b, subplot on 30×). At low (10× and 20×) coverage, PASS performed reasonably well. MuSE could not identify a cutoff comparable to the MuTect default cutoff for 20× coverage, VAF of 0.1 and 0.05. Tier–5 was helpful to improve sensitivity while maintaining a low FPR at low coverage and low VAF. Looking across the data sets with varied coverage but fixed low VAF (VAF=0.05), we observe that MuSE achieved higher sensitivity than MuTect at the same level of specificity. Therefore, MuSE will be helpful for calling subclonal mutations in studies of the heterogeneity and subclonal evolution of tumors. Although MuSE demonstrated better accuracy than MuTect using the virtual-tumor benchmarking data, the two callers generated intersecting sets (Supplementary Fig. 1), which provides a conspicuous demonstration of the importance and necessity of using multiple callers in somatic variant detection.

### Real data

We evaluated MuSE using multiple real WES and WGS data sets and compared MuSE with other calling pipelines (anonymous, see pipeline names in supplementary material 2). Specifically we focused on comparing with Caller–A, which is one of the best-in-breed mutation callers based on the ICGC-TCGA DREAM Mutation Calling challenge. With TCGA and ICGC samples, we used calls that were prepared and provided by the corresponding institutes where individual calling pipelines were ran. We first tested the performance of MuSE using data from 91 tumor-normal paired WES samples (3.21 Gbp) from patients with adrenocortical carcinoma (ACC; Fig. 3a). Taking into account the tier-based distribution of MuSE calls (Supplementary Table 2), we computed the validation rates of MuSE total calls and unique calls, and obtained 84.50% and 26.34%, respectively. We repeated a similar calculation for Caller–A, which gave the respective validation rates of total calls and unique calls as 87.39% and 24.79%. Considering that the validation rate could not measure sensitivity, we extracted the multi-center somatic variant calls from the TCGA mutation annotation format (MAF) file, made an artificial truth set by taking calls that were shared by at least three callers, and computed a sensitivity of 98.71% for MuSE and a sensitivity of 95.10% for Caller–A (Fig. 3b). Moreover, MuSE missed only 7 calls that were captured by the other four callers, comparing with 66, 36, 807 and 1,626 missed calls from Caller–A, Caller–1, Caller–2 and Caller–3, respectively. As an alternative to the deep sequencing validation on a small set of positions, we regarded all calls outside of the artificial truth set as false positives to calculate positive predictive values (PPVs). In agreement with previous findings of the validation rates, Caller–A benefited from its low number of unique calls and obtained the second best PPV, which in turn helped Caller–A acquire a better F_1_ score [12]. However, using the F2 score, which placed a relatively higher weight on sensitivity, we demonstrated the good performance of MuSE (F_2_;= 0.9366). When we used more stringent tiers, we obtained a smaller number of MuSE unique calls, changing from 2,152 to 378, without losing much sensitivity, i.e., the number of missed calls that were shared by the other four callers increased from 7 to 14 (Supplementary Fig. 2).

**Figure 3.**
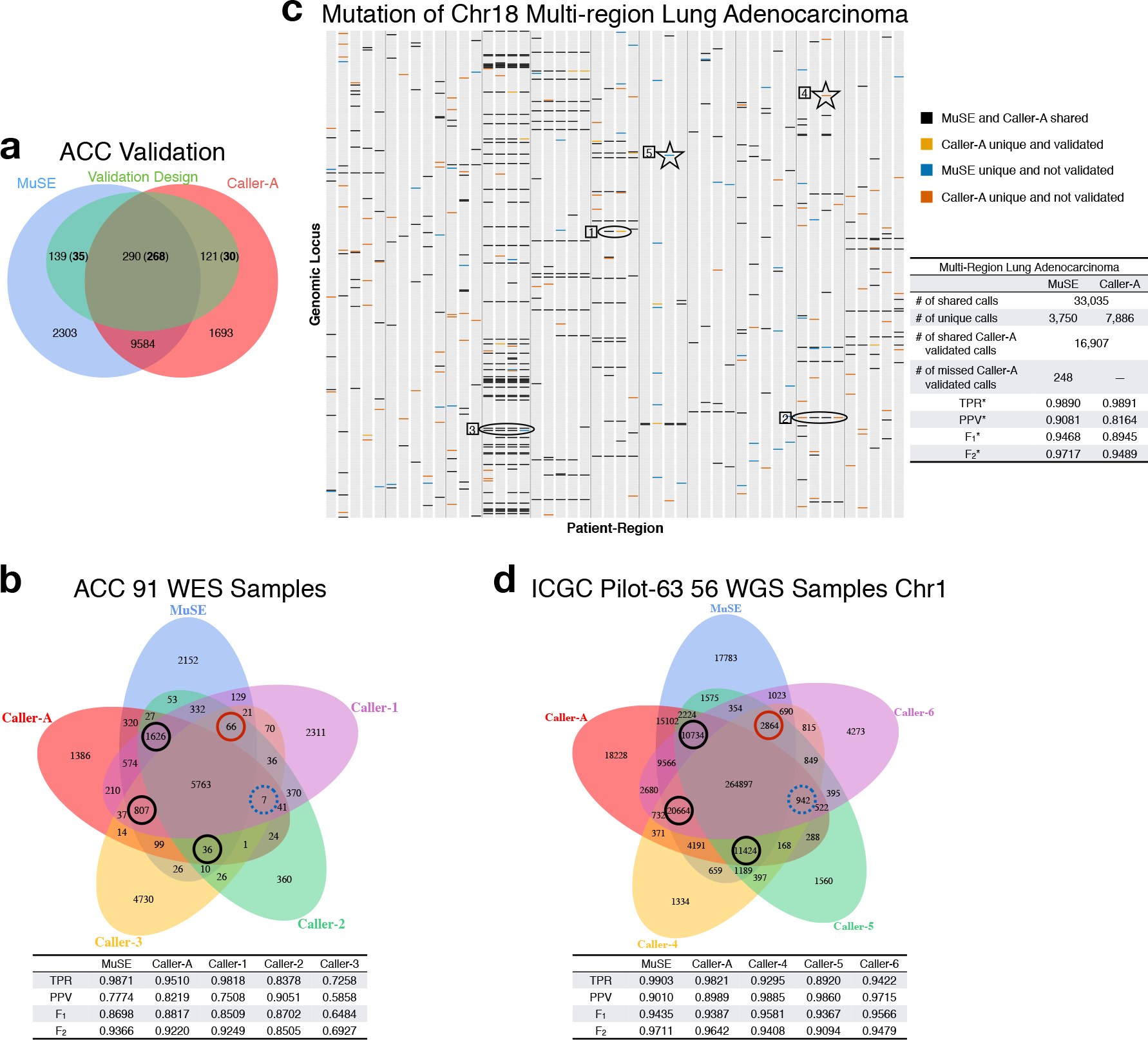
MuSE performance in two WES data sets and one WGS data set. (a) Venn diagram of MuSE and Caller–A calls from 91 pairs of ACC WES samples. The calls are overlaid with 550 positions that were selected for deep sequencing validation. The numbers of validated calls are shown in boldface. For selected MuSE unique, MuSE Caller–A shared and Caller–A unique calls, 35 out of 139, 268 out of 290 and 30 out of 121 are validated, respectively. (b) Venn diagram of calls from five different callers using the same ACC data. All the calls except those of MuSE are extracted from TCGA mutation annotation format (MAF) file. The circles label the numbers of calls missed by one caller but captured by the other four callers. The blue dotted circle denotes the number of calls missed by MuSE, and the red solid circle indicates the number of calls missed by Caller–A. TPR, PPV, F_1_ and F_2_ scores are calculated and listed below the Venn diagram. The truth set is defined as calls shared by at least three callers. (c) Mutation plot and summary table of MuSE and Caller–A calls from 48 pairs of multi-region lung adenocarcinoma WES samples. Each gray column represents a sample. MuSE and Caller–A share 33,035 calls and possess 3,750 and 7,886 unique calls, respectively. Only calls from Caller–A were further validated. MuSE confirms 16,907 and misses 248 Caller–A validated calls. Calls from chromosome 18 are shown in the mutation plot to illustrate how the artificial truth set and false positives are defined. The vertical gray lines separate 11 patients who have samples from 3 to 5 regions of one tumor. The numbered shapes combined with different call types are examples for defining the artificial truth set as positions that fall into any of the three categories: shared or validated (oval–1), called in all regions by including Caller–A unique (oval–2), called in all regions by including MuSE unique (oval–3), and false positives: unique and single calls (star 4 and star 5). Correspondingly, the TPR^*^, PPV^*^, 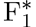 and 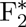 scores are calculated and listed besides the mutation plot. (d) Venn diagram of calls from five different callers using 56 pairs of ICGC Pilot–63 WGS samples on chromosome 1. The circles label the number of calls missed by one caller but captured by the other four callers. The blue dotted circle denotes the number of calls missed by MuSE, and the red solid circle indicates the number of calls missed by Caller–A. TPR, PPV, F_1_ and F_2_ scores are calculated and listed below the Venn diagram.

We then applied MuSE to WES data from 48 multi-region tumor-normal paired samples (2.46 Gbp) from 11 patients with lung adenocarcinoma, which provided 17,155 deep sequencing validated calls that were originally selected from all calls made by Caller–A [13]. MuSE confirmed 16,907 and missed 248 Caller–A validated calls, a sensitivity of 98.55%, given 3,750 unique calls compared with 7,886 unique calls from Caller–A. In contrast to the ACC data, this validated data set could not provide unbiased evaluation of the two callers. However, the multi-region design of this data set was unique. We therefore built our artificial truth set by taking all validated calls (Fig. 3c; orange in oval 1), all shared calls (Fig. 3c; black in ovals 1) and all trunk mutation calls that occurred at the same genomic locus in all tumor regions of one patient (Fig. 3c; oval 2 and 3). This design allowed us to consider unique and unvalidated calls from each caller as true positives when they appeared as trunk mutations (Fig. 3c; red in oval 2 and blue in oval 3). We regarded all other calls that were subclonal as false positives (Fig. 3c; red in five-pointed star 4 and blue in five-pointed star 5). The 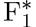 (0.9468) and 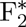 (0.9717) scores acquired by MuSE were higher than those of Caller–A.

We further compared MuSE with other callers using 56 pairs of ICGC Pilot-63 WGS samples on chromosome 1 [14.0 Gbp; 14]. We downloaded the related somatic VCF files that were generated by multiple callers from the ICGC Pilot–63 study. In accordance with the ACC multi-caller result, MuSE missed 942 calls that were captured by the other four callers, which was the least number of missed calls and therefore indicated the highest sensitivity among all five callers (Fig. 3d). Caller–4 and Caller–6 gave the best and the second best F_1_ scores due to their high PPVs (Fig. 3d). Caller–5, which had low sensitivity, could not achieve a better F_1_ score, although its PPV was higher than that of Caller–6. The F_1_ score of MuSE was higher than those of Caller–A and Caller–5, but could not compete with those of Caller–4 and Caller–6. However, considering that Caller–4, Caller–5 and Caller–6 respectively missed 10,734, 20,664 and 11,424 calls that were shared by the other four callers, the loss of sensitivity as a tradeoff for greater specificity may raise concerns. Among all five callers, MuSE had the best F_2_ score, emphasizing the importance of sensitivity.

## Conclusions

In summary, we present a somatic point mutation caller, MuSE. We design MuSE as an automatic approach with two steps. The first step, ‘MuSE call’, implements the heuristic pre-filters and uses the Markov substitution model to describe the evolution of the reference allele to the allelic composition of the matched tumor and normal tissue at each genomic locus, which provides the summary statistics *π*_somatic_. The *π*_somatic,tumor_ associated ROC curve is shown to stand above that from Caller–A, suggesting the good ability of discriminating mutations from references of the MuSE pipeline. The second step, ‘MuSE sump’, identifies tier-based cutoffs on *π*_somatic,tumor_. We build a sample-specific error model to account for tumor heterogeneity, and to identify cutoffs that are unique to each sample, achieving high accuracy in mutation calling. With the two steps, we aim at mitigating users’ curation of output. We provide 5 tiers. In experience, we suggest using calls up to Tier–4 for WES data, and calls up to Tier–5 for WGS data. These suggested cutoffs are derived based on our observation of real data and serve the goal of maximizing sensitivity and maintaining a good specificity. Typically, ‘MuSE call’ step takes ~4 hours to process a tumor-normal paired WGS sample with 30 − 60× coverage when the WGS data is divided into ~50 equal-sized blocks and each block is assigned with 1 CPU core and 2 GB memory, and ‘MuSE sump’ step requires ~1 hour for WGS data given 1 CPU core and 4 GB memory.

We demonstrate the reliable performance of MuSE using both synthetic and real data, such as the ICGC-TCGA DREAM Mutation Calling challenge WGS data, the virtual-tumor benchmarking approach, TCGA ACC WES data, the multi-region lung adenocarcinoma WES data and the ICGC PanCancer Pilot–63 WGS data. We demonstrate the superior sensitivity of MuSE, especially to low VAF mutations, and its capacity to identify an appropriate balance of sensitivity and specificity in each sample with varying levels of heterogeneity. This feature is essential for downstream analyses, such as finding tumor subclonal structures and understanding the evolution of tumors, a broad interest in the cancer community and beyond. So far, we have found substantially more subclones using MuSE calls (up to Tier–5) than using calls from other callers in ICGC PanCancer Analysis of Whole Genomes [data not shown; 15].

Copy number aberration (CNA), tumor purity and tumor subclonality commonly exist in our data, both synthetic and real. All influences of CNA, tumor purity and tumor subclonality on the mutant chromosome content of a tumor reduce to the same question of VAF, and the mechanism of creating or changing the VAF is not as important as the VAF itself in terms of somatic mutation calling. Therefore, we use the F81 Markov substitution model to capture the VAF dynamics at each locus. Our *π*_somatic,tumor_ is directly related with the configuration of local copy number variation, purity and subclonality of the position. Our sample-specific error model aims to deconvolute two log (2*π*_somatic,tumor_) distributions, one from true mutations and one from reference positions. When there are multiple peaks in each mode as often observed in real data, our assumption that the signal and noise can be separated by finding two major modes is supported by the high sensitivity and specificity of MuSE calls in the validation data.

We considered two aspects when using the F81 model: (1) the number of free parameters in the model should remain small to allow for higher accuracy in estimation for each position; and (2) the F81 model can be extended to take into account mutational contexts, which will be our future work. One potential benefit of considering the mutational contexts is to further reduce false positives. We accessed MuSE calls in annotated CpG islands (UCSC Genome Browser CpG Island Annotation Track) using the TCGA WES data from ACC. The validation rate of MuSE total calls decreased from 0. 8450 to 0.7245, and the validation rate of MuSE calls shared with Caller–A decreased from 0.9889 to 0.8829.

We will further validate MuSE through participation in the ICGC-TCGA DREAM Mutation Calling challenge and the ICGC Pilot–63 study, both of which have promised independent experimental validations. We have also applied MuSE to analyze the WES data of chromophobe renal cell carcinoma [KICH; 16] and liver hepatocellular carcinoma (LIHC), which are part of TCGA project. The corresponding calls have been made available to TCGA community. MuSE is being used by two new ongoing consortium projects: TCGA PanCanAtlas and ICGC PanCancer Analysis of Whole Genomes that includes WGS data from over 2,800 pairs of tumor and matched-normal samples.

Despite the satisfactory performance of MuSE, we contend that there is no comprehensive caller that can replace all the others; each caller has strengths and unique attributes. We support the trend to incorporate call sets from multiple callers in future NGS analyses, for example, using SomaticSeq [17]. Due to its ensemble nature, SomaticSeq relies on the performance of its callers, and is bounded by the best sensitivity among individual callers [17]. Therefore when MuSE is included as one of the callers to be v—integrated, we expect SomaticSeq to generate results that are more accurate than it can do currently. We welcome the usage of other post filtering methods on MuSE calls, for instance, panel of normal samples, when data from the appropriate control samples are available. Our method can be extended for calling bi-nucleotide, triplet or small insertion-deletion variants by modifying the F81 Markov substitution model.

MuSE is freely available for noncommercial use [18].

## Methods

### Data sets

(1) We downloaded the WGS BAM files of NA12878 and NA12981 [19], and followed the procedures described in Cibulskis *et al.* [5] to simulate the virtual-tumor benchmarking data set.
(2) We downloaded the ICGC-TCGA DREAM Mutation Calling challenge IS1, IS2 and IS3 WGS BAM files and VCF files of simulated truths [20].
(3) According to TCGA cancer types, we downloaded the tumor-normal paired WGS BAM files of 3 patients in LUAD, LUSC and STAD, and the tumor-normal paired WES BAM files of 91 patients in ACC from UCSC Cancer Genomics Hub (CGHub).
(4) We obtained the tumor-normal paired WES BAM files of 11 lung adenocarcinoma patients, including 48 multi-regions that were collected and generated at MD Anderson Cancer Center [13].
(5) We obtained somatic SNV VCF files of 56 samples that were generated by multiple callers from the ICGC Pilot-63 study [21].

### BAM preparation

All the sequence reads were aligned against the hg19 reference genome using the Burrows-Wheeler alignment tool (BWA), with either the backtrack or the maximal exact matches (MEM) algorithm [22]. In addition, data sets (3), (4) and (5) were processed by following the Genome Analysis Toolkit (GATK) Best Practices [23, 24, 25] that include marking duplicates, realigning the paired tumor-normal BAMs jointly and recalibrating base quality scores.

### Variant heuristic pre-filters

In order to detect context-based sequencing artifacts, remove potential false positives and accelerate the computing speed, we apply heuristic pre-filters to every genomic locus in advance of variant detection.

(1) *Neighboring to indels*: No less than 3 insertions or 3 deletions are observed in an 11-base window centered on the locus.
(2) *Variant in matched-normal*: The candidate variant allele is observed no less than twice or its variant allele fraction is no less than 3% in the matched-normal data, and moreover, the sum of the variant allele’s base quality scores is more than 20. However, this genomic locus is kept if the candidate variant allele turns out to be the germ-line variant in the matched-normal data and the second variant allele is rejected by the above test.
(3) *Minimum variant allele fraction*: The candidate variant allele fraction in the tumor data is smaller than 0.005.
(4) *Strand bias*: The p-value that is computed from Fisher’s exact test using tumor allele count data comparing sense and antisense strands is less than or equal to 1e–5.
(5) *Dubious mapping quality*: The average mapping quality score of reads that carry a candidate variant allele is less than or equal to 10.
(6) *Read-end cluster*: For each read that has the candidate variant allele, we record the smallest distance that can be from the current genomic locus to either the left end or the right end of the read alignment. We disregard the current genomic locus if the median of all these distances is less than or equal to 10 and the median absolute deviation is less than or equal to 3.
(7) *Confident variant*: We require there is at least one variant read that meets the following criteria: (i) the read and its mate are mapped in a proper pair; (ii) its mapping quality score is no less than 30; and (iii) the base quality score of its candidate variant allele is greater than or equal to 25.

### Variant detection

For each genomic locus, we denote the base of read *r* (*r* = 1 … *N*) that covers the locus as *b_r_*, where *r* ∈ {1…*N*} and *N* is the depth of the locus. By knowing the associated Phred quality score *q_r_* of *b_r_*, we denote the probability of *b_r_* being the four different alleles (A, C, G, T) as

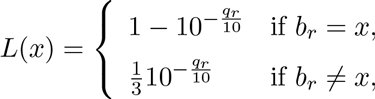

where *x* ∈ {A, C, G, T}. We use a continuous-time Markov chain to describe the DNA evolution from the reference allele *R* to the allelic composition ***b*** = (*b_r_*: *r* ∈ {1…*N*}) at each locus, namely, the F81 Markov substitution model [9]. The F81 model can be expressed using a 4-state × 4-state instantaneous rate matrix, *Q*,

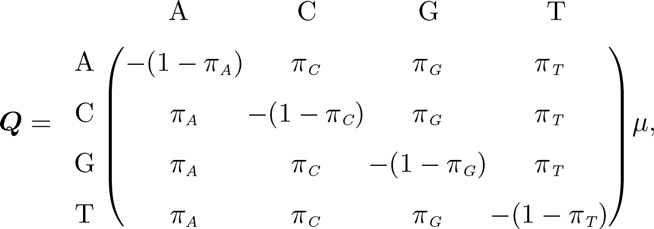

where each entry represents the changing rate from allele *i* to allele *j* in an infinitesimal time *dt*, *μ* stands for the mean instantaneous substitution rate, and *π_A_*, *π_C_*, *πn_a_*, *π_T_* are the equilibrium allele frequencies. The transition matrix that consists of the probabilities of change between any two states in time *t* can be calculated from the exponential of the instantaneous rate matrix, **P**(*t*) = *e^Qt^*. Specifically,

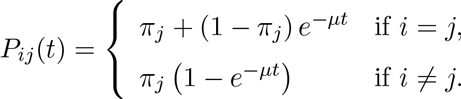

Because of the confounding nature of the *μt* product, it is ordinary to rescale the instantaneous rate matrix so that the mean substitution rate at equilibrium is 1, and replace *t* with the evolutionary distance *ν* that represents the expected number of substitutions per base. Consequently, the transition matrix of the F81 model is altered as

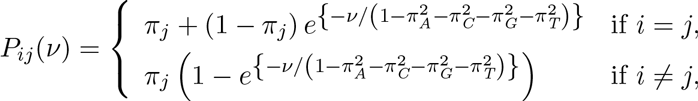

and the likelihood function *f* (***b***, *R*|*π*, *ν*) can be expressed as

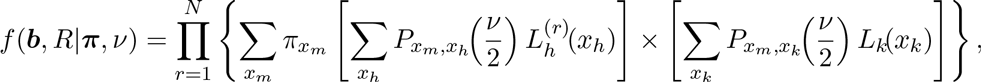

where (1) *x_m_*, *x_h_*, *x_k_* ( {A, C, G, T}; (2) *ν* connects the reference allele R and the allelic composition ***b***; (3) *h* and *k* denote the ***b*** and *R* tips of *ν* respectively; (4) *m* denotes the middle point of *h* and *k* so that the evolutionary distance *ν* from *m* to *h* is equal to the distance from *m* to *k*, i.e., *ν*/2. Because of the time reversible characteristic of F81 model, *m* can be any point along the evolutionary distance *ν* that connects the *h* and *k* tips without affecting the final result. We set *m* as the mid-point for the purpose of calculation convenience; (5) *L_k_*(*x_k_*) = 1 if *x_k_* = *R*, and *L_k_*(*x_k_*) = 0 otherwise; and (6) all the other notations are the same as above.

We obtain the joint posterior probabilities of ***π*** and *ν*, *f* (***π***, *ν*|***b***, *R*), by setting the priors of ***π*** and *ν* to be the Dirichlet distribution Dir(1,1,1,1) and the exponential distribution Exp(1000), respectively. In practice, we employ the Broyden-Fletcher-Goldfarb-Shanno algorithm and Brent’s algorithm to search for the maximum a posteriori (MAP) estimates of ***π*** and *ν* instead of exploring the full posterior distribution.

We apply the above method to both loci of the tumor-normal paired sequencing data, and obtain the *π*_somatic,tumor_ and the *π*_somatic,norma_i estimates accordingly. We designate the non-reference and non-germline allele that has the largest *π* as the somatic variant allele. The somatic variant allele should pass all the pre-filtering examinations.

### Post-filtering criteria

After we obtain the *π*_somatic,tumor_ and *π*_somatic,normal_, we require that: (1) the minimum coverage of tumor and matched-normal data is 8 at given genomic loci; and (2) the ratio 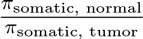 is less than or equal to 0.05, which tolerates the contamination of matched-normal data with tumor data in a reasonable amount and dynamically changes the constraint on matched-normal data.

### Sample-specific error model

We provide two options for building the sample-specific error model. One is applicable to WES data, and the other to WGS data. By plotting the densities of log (2*π*_somatic,tumor_) from MuSE on all positions (see Supplementary Fig. 3), we observed, (1) the density of log-transformed *π*_somatic,tumor_ showed a bi-modal behavior that could be approximated using a Gaussian mixture distribution; (2) the true positives (red) and reference positions (blue) correspond to each of the mode so that a cutoff can be identified to separate the two types of calls; (3) as expected, the separation of two modes becomes easier at higher coverage and higher variant allele fraction (VAF). For most WES data, there are not enough true mutations that can form a detectable second mode as compare to the reference positions. As *π*_somatic,tumor_ provides a good ranking of true versus false mutations, we fit a Beta distribution on the *π*_somatic,tumor_ in this case and call mutations as the extreme and rare events on the right tail of the fitted distribution. For the WGS data, we transform all post-filtered *π*_somatic,tumor_ to the logarithmic scale and then fit a two-component Gaussian mixture distribution on it. Given the means *μ*_1_ and *μ*_2_, standard deviations *σ*_1_ and *σ*_2_, and weights *p* and 1 − *p* of the two Gaussian distributions that are estimated using the expectation-maximization algorithm, we first calculate the cutoff that minimizes the misclassification, the sum of the false positive probability and the false negative probability,

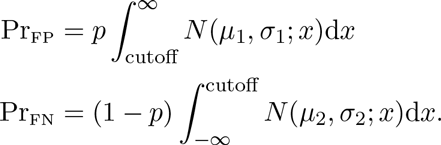

If the cutoff is larger than 0.01, we consider it as PASS and 0.01 as Tier–1, or vice versa. We take the top 0.1, 0.5 and 1 percentiles of the Tier–1 truncated Gaussian distribution as Tier–2, Tier–3 and Tier–4, respectively. For the WES data, we build the sample-specific error model upon post-filtered *π*_somatic,tumor_ that are within the interval (0.0025, 0.01). We first rescale all selected *π*_somatic,tumor_ to the range (0, 1), and then fit a beta distribution on them. We report 0.01 as PASS, and cutoffs that are transformed from the top 0.1, 0.5, 1 and 2 percentiles of the beta distribution as Tier–1, Tier–2, Tier–3 and Tier–4, respectively.

### Sensitivity and specificity

For the virtual-tumor benchmarking data, we measured sensitivity and specificity by applying MuSE and MuTect [5] to the combination of 24 spike-in BAMs (4 different variant allele fractions × 6 distinct depths) with the same depth non-spike-in WGS BAMs. The matched-normal WGS BAM was fixed at 30× depth. We considered any missed calls from our *in silico* spike-in ground truth as false negatives, and any calls from the non-spike-in WGS BAMs as false positives. The denominator for the FPR calculation is the total length of the hg19 reference genome from chromosome 1 to chromosome X.

For the DREAM challenge IS1, IS2 and IS3 data, we took the organizer provided script and the truth VCF files to compute sensitivity and specificity [20]. We extracted sensitivity and specificity of SomaticSniper, Strelka and VarScan2 from the DREAM challenge leader-boards. The denominator for the FPR calculation is the total length of the hg19 reference genome from chromosome 1 to chromosome X.

For the multi-region lung adenocarcinoma data, we calculated sensitivity and the positive predictive value (PPV) based on an artificial truth set for the reason that the known validation set was extracted and compiled from the papers supplementary document and was biased toward Caller–A. The artificial truth set included shared calls (Figure 3c; black in ovals 1, 2 and 3), validated calls (Figure 3c; orange in oval 1) and unique-not-validated calls that helped the recognition of trunk mutations (Figure 3c; red in oval 2 and blue in oval 3). Here, a trunk mutation was a somatic variant call that all tumor regions of one patient had at the same genomic locus. All the other calls were considered as false positives (Figure 3c; red in five-pointed star 4 and blue in five-pointed star 5). We evaluated accuracy using the F_1_ and F_2_ scores that were defined as

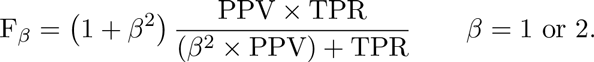

To compare the performance of multiple callers in the ACC WES data and the ICGC Pilot–63 WGS data, we also made the artificial truth sets by taking calls that were shared by at least 3 callers, and computed sensitivity. We regarded other calls as false positives to calculate PPVs. We calculated the F_1_ and F_2_ scores by following the same equation above.

## Validation

To validate variants identified by MuSE and Caller–A in the ACC data, we selected 550 patient-specific positions and designed NimbleGen probes correspondingly for the purpose of targeted capture enrichment and deep sequencing. Paired-end Illumina re-sequencing was carried out to an average sequencing depth at 1500×. After mapping the reads against the hg19 reference genome using BWA, we considered a somatic variant as validated if its p-value calculated from Fisher’s exact test comparing the tumor and matched-normal samples was not larger than 0.05. The validation rates of MuSE and Caller-A were calculated as

validation rate of MuSE unique calls

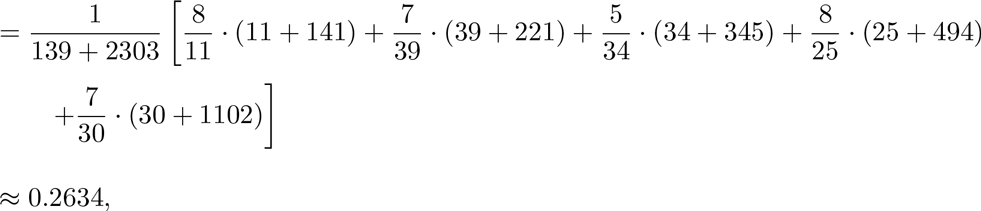

validation rate of MuSE shared calls

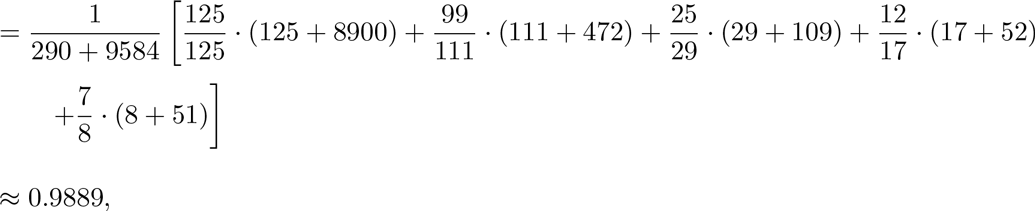

validation rate of MuSE total calls

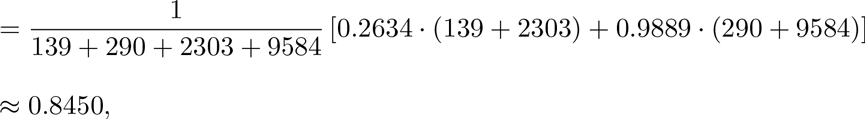

validation rate of Caller-A unique calls

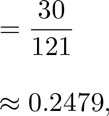

validation rate of Caller-A total calls

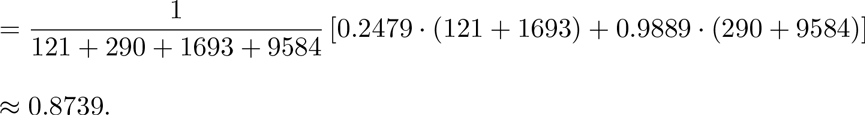

## Authors’ contributions

Y.F. and W.W. designed the study. Y.F. designed and developed MuSE and performed the analysis. L.X., D.S.T.H., J.Z. and J.Z. assisted in the analysis. Y.F., D.A.W. and W.W. wrote the manuscript. P.A.F. critically reviewed the manuscript. W.W. led the project. All authors discussed the results. All authors read and approved the final manuscript.

## Acknowledgements

We thank the ICGC PCAWG for use of the ICGC Pilot–63 data and Kyle Ellrott from UCSC for generating the MuSE calls presented in Figure 3d. We thank Dr. John N. Weinstein from MDACC and Dr. Richard A. Gibbs from BCM HGSC. Y.F. is partially supported by a training fellowship from the Keck Center of the Gulf Coast Consortia for the Computational Cancer Biology Training Program; the Cancer Prevention and Research Institute of Texas (CPRIT) RP140113, PI-Rathindra Bose; and the National Institutes of Health/National Cancer Institute through grant U24 CA143883 02S2 (to J.N.W.) and the Integrative Pipeline for Analysis & Translational Application of TCGA Data, grant 5U24CA143883–04 (to J.N.W.). W.W. is supported in part by the Cancer Prevention Research Institute of Texas through grant number RP130090, and by NCI through grant numbers 1R01CA174206-01, 1R01CA183793-01 and P30 CA016672. Y.F. and W.W. are supported in part by U.S. National Cancer Institute (NCI; MD Anderson TCGA Genome Data Analysis Center) through grant numbers CA143883 and CA083639. J.Z. and P.A.F. are supported by the Cancer Prevention and Research Institute of Texas through grant number R1205 01, the UT Systems Stars Award (PS100149), the Welch Foundation Robert A. Welch Distinguished University Chair Award (G–0040), MD Anderson Physician Scientist Award, and C.G. Johnson Advanced Scholar Award.

## SUPPLEMENTARY TABLES

Supplementary Table 1: Number of positive spike-in conditions for the virtual-tumor benchmarking approach. Tumor data coverage varies from 10x to 60x and spike-in VAFs are 0.05, 0.1, 0.2 and 0.4.

Supplementary Table 1: Number of positive spike-in conditions for the virtual-tumor benchmarking approach. Tumor data coverage varies from 10 × to 60 × and spike-in VAFs are 0.05, 0.1, 0.2 and 0.4.
| Spike-in VAF | Tumor Data Coverage |  |  |  |  |  |
| --- | --- | --- | --- | --- | --- | --- |
|  | 10x | 20x | 30x | 40x | 50x | 60x |
| 0.05 | 608,275 | 608,403 | 608,438 | 608,469 | 608,482 | 608,486 |
| 0.1 | 608,274 | 608,403 | 608,431 | 608,444 | 608,446 | 608,400 |
| 0.2 | 608,274 | 608,374 | 608,328 | 608,172 | 607,497 | 605,323 |
| 0.4 | 608,251 | 608,088 | 605,272 | 586,219 | 516,971 | 379,905 |

Supplementary Table 2: Allocation of MuSE and Call–A calls for the ACC data. Calls are categorized by whether they are selected for deep sequencing validation and whether they are unique for one caller or shared by two callers. The number of validated calls is shown in parentheses.

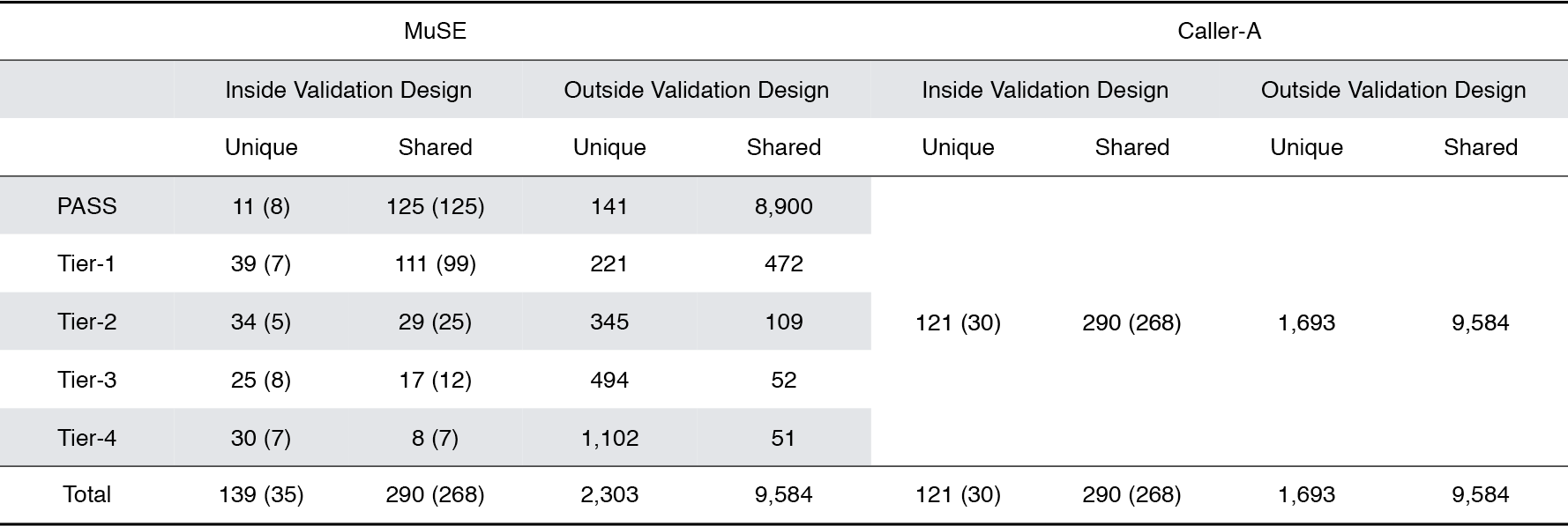

## SUPPLEMENTARY FIGURES

**Venn Diagrams of MuSE and MuTect Calls Using the Virtual-tumor Benchmarking Data**Supplementary Figure 1: Venn diagrams of MuSE and MuTect calls compared with the truth across ball combinations of tumor data coverage and VAFs for the virtual-tumor benchmarking data.

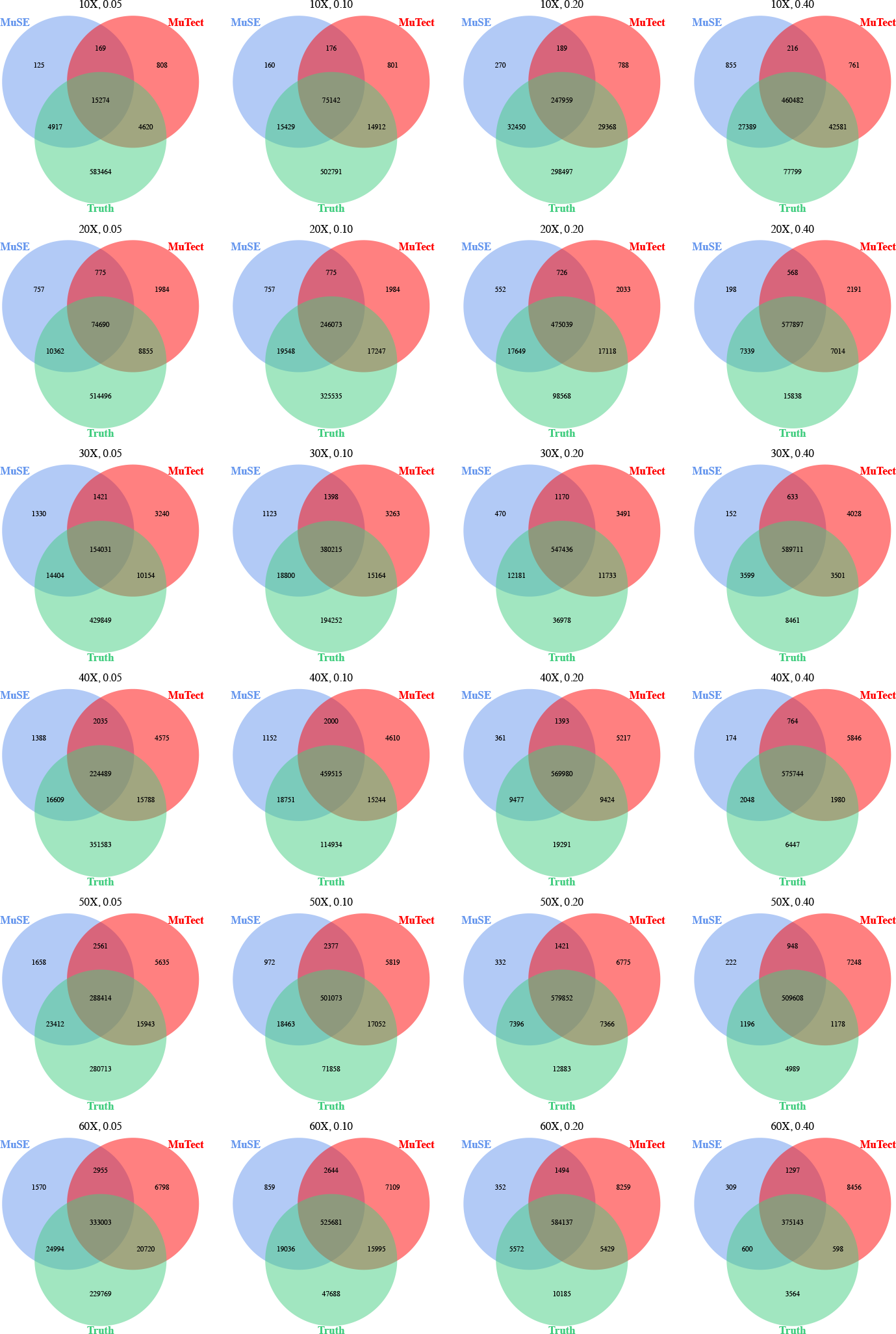

**Venn Diagrams of Calls from Five Different Callers Using the ACC Data** Supplementary Figure 2: Venn diagrams of calls from five different callers using the ACC data. All the calls except those of MuSE are extracted from TCGA mutation annotation format (MAF) file. MuSE calls from different cutoffs (PASS only, up to Tier–1, up to Tier–2 and up to Tier–3) are used. The blue circles label the number of MuSE unique calls and the number of calls missed by MuSE but captured by the other four callers. The red circles have the same meaning but apply to Caller–A.

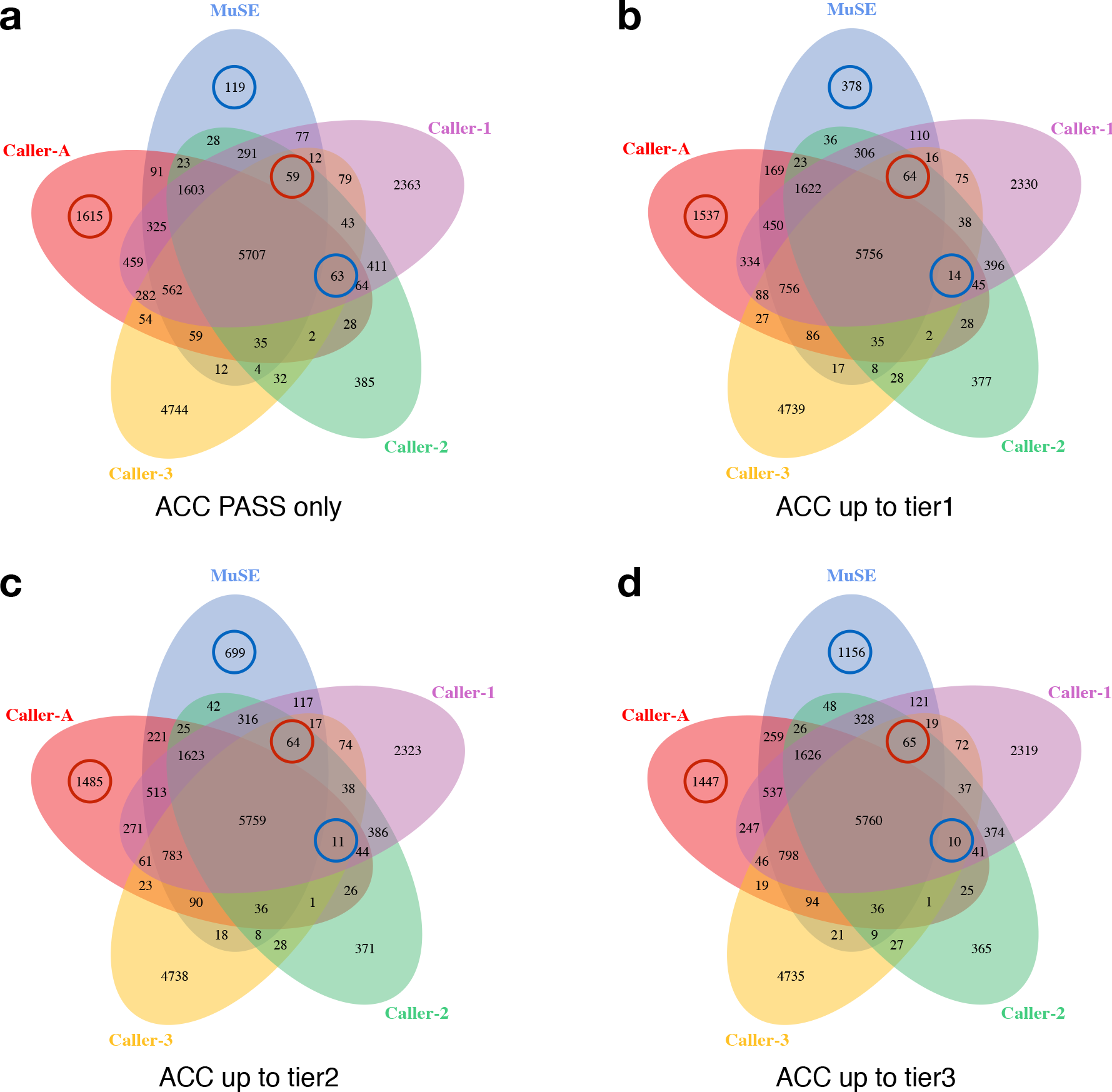

**Density Distributions of log (2*π*_somatic;tumor_) from MuSE on the Virtual-tumor Benchmarking Data** Supplementary Figure 3: Density distributions of log (2*π*_somatic,tumor_) from MuSE on the virtual-tumor benchmarking data. Rows correspond to different coverage depth, from 10× to 60×, at an increment of 10×. Every two columns correspond to a different scenario for VAF, at 0.05, 0.1, 0.2 and 0.4, from left to right. In each pair of columns, the panel on the left shows the density of true mutations (in red) and that of the remaining reference positions (in blue), respectively. The panel on the right shows the density of all positions together without the knowledge of mutation status.

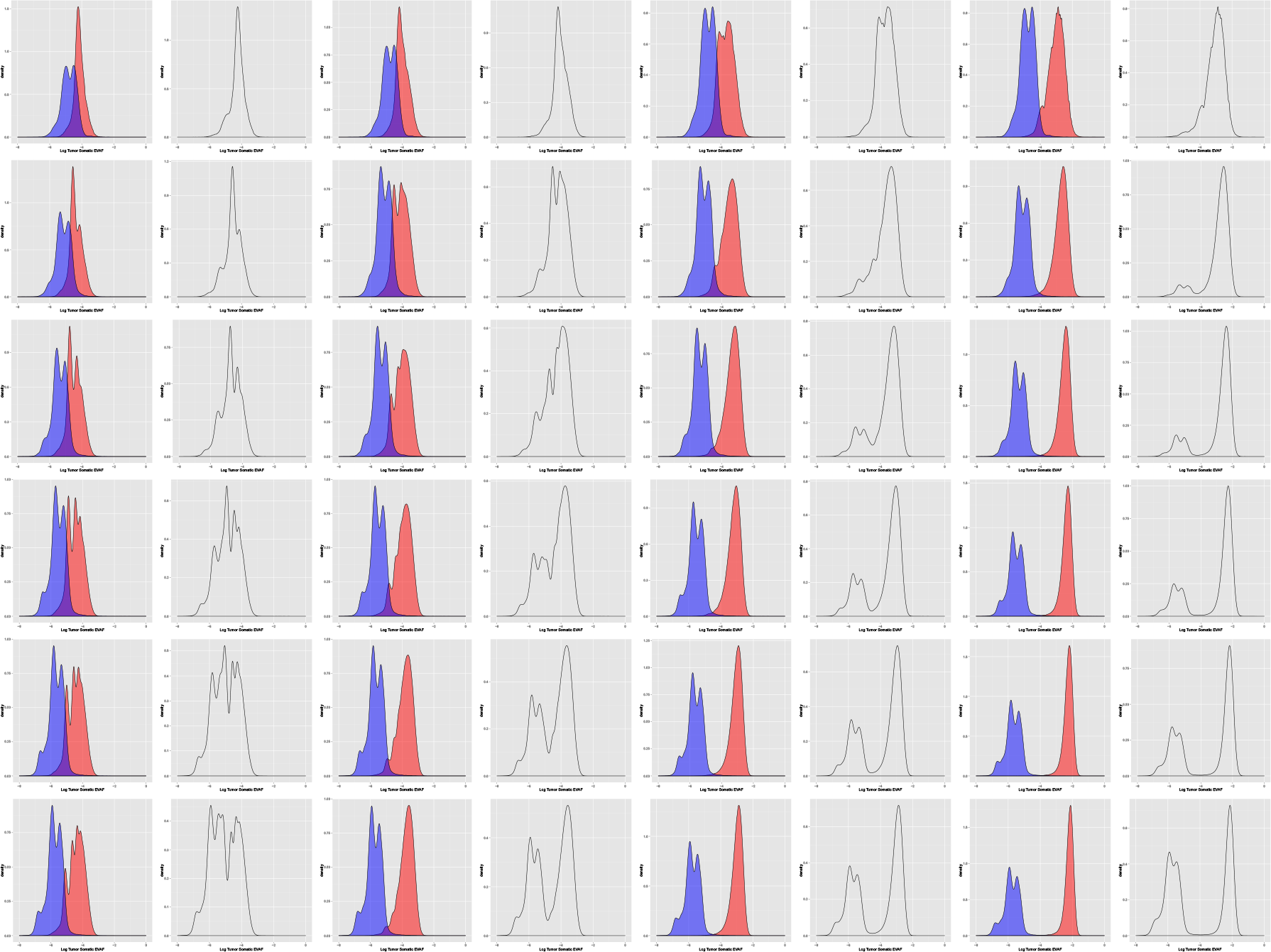

